# *DeepRetroMoCo:* Deep neural network-based Retrospective Motion Correction Algorithm for Spinal Cord functional MRI

**DOI:** 10.1101/2022.09.06.506787

**Authors:** Mahdi Mobarak-Abadi, Ahmad Mahmoudi-Aznave, Hamed Dehghani, Mojtaba Zarei, Shahabeddin Vahdat, Julien Doyon, Ali Khatibi

## Abstract

There are unique challenges in the preprocessing of spinal cord fMRI data, particularly voluntary or involuntary movement artifacts during image acquisition. Despite advances in data processing techniques for movement detection and correction, there are challenges in extrapolating motion correction algorithm developments in the brain cortex to the brainstem and spinal cord. We trained a Deep Learning-based convolutional neural network (CNN) via an unsupervised learning algorithm, called DeepRetroMoCo, to detect and correct motions in axial T2*-weighted spinal cord data. Spinal cord fMRI data from 27 participants were used for training of the network (135 runs for training and 81 runs for testing). We used average temporal signal-to-noise-ratio (tSNR) and Delta Variation Signal (DVARS) of raw and motion-corrected images to compare the outcome of DeepRetroMoco with sct_fmri_moco implemented in the spinal cord toolbox. The average tSNR in the cervical cord was significantly higher when DeepRetroMoco was used for motion correction compared to sct_fmri_moco method. Average DVARS was lower in images corrected by DeepRetroMoco than those corrected by sct_fmri_moco. The average processing time for DeepRetroMoco was also significantly shorter than sct_fmri_moco. Our results suggest that DeepRetroMoCo improves motion correction procedures in fMRI data acquired from the cervical spinal cord.

## 1. Introduction

Spinal cord functional magnetic resonance imaging (fMRI) has become increasingly popular for exploring intrinsic neural networks and their role in pain modulation, motor learning and sexual arousal (Alexander et al., 2016; Kinany et al., 2019). There are unique challenges in data acquisition and preprocessing, such as relatively small cross-sectional dimension, the variable articulated structure of the spine between individuals, low signal intensity in standard gradient-echo echo-planar T2^∗^-weighted fMRI and voluntary (bulk motion) or involuntary (fluctuation of cerebrospinal fluid due to respiration and heartbeat) movements during image acquisition (Dehghani, Oghabian, Batouli, Arab Kheradmand, & Khatibi, 2020; Kinany et al., 2022; Powers, Ioachim, & Stroman, 2018). Spinal cord motions can cause signal alterations across volumes, which decrease the temporal stability of the signal and ultimately increase false positive and negative discovery rates (Cohen-Adad, Piche, Rainville, Benali, & Rossignol, 2007; Dehghani, Weber, Batouli, Oghabian, & Khatibi, 2020; Stroman, Figley, & Cahill, 2008).

Despite advances in fMRI motion correction, there are problems in extrapolating the motion correction algorithm developments in the brain to the brainstem and spinal cord. In brain fMRI, we generally utilize six degrees of freedom rigid-body registration of a single volume to a reference, which can be a preselected volume or an average volume (Maknojia, Churchill, Schweizer, & Graham, 2019; Oakes et al., 2005). This method is non-robust and insufficient for spinal cord fMRI preprocessing due to the non-rigid motion of the spinal column and physiological motion from swallowing and the respiratory cycle (Dehghani, Oghabian, et al., 2020; Fratini, Moraschi, Maraviglia, & Giove, 2014). Along with the release of the Spinal Cord Toolbox (SCT), sct_fmri_moco was introduced for motion correction in the spinal cord (De Leener et al., 2017). The basis of sct_fmri_moco is slice-by-slice regularized registration for spinal cord algorithm (SliceReg) that estimates slice-by-slice translations of axial slices while ensuring regularization constraints along the z-axis (Paquin et al., 2018).

In the past few years, we have seen an interest in the application of artificial intelligence in medical image processing (Anaya-Isaza, Mera-Jiménez, & Zequera-Diaz, 2021; Varoquaux & Cheplygina, 2022; Wen et al., 2018). In spinal cord imaging, deep learning has been used for the segmentation of the spinal cord and CSF in structural T1 and T2 weighted images. DeepSeg as a fully automated framework based on convolutional neural networks (CNNs), is proposed to apply spinal cord morphometry for segmenting the spinal cord, as part of SCT (Gros et al., 2019; Perone, Calabrese, & Cohen-Adad, 2018; Prados et al., 2017). More recently, the K-means clustering algorithm has been used for the segmental spinal cord in the thoracolumbar region (Sabaghian, Dehghani, Batouli, Khatibi, & Oghabian, 2020). A robust and automated CNN model with two temporal convolutional layers is introduced for motion correction in brain fMRI, and the following regression employs derived motion regressors. (Yang, Zhuang, Sreenivasan, Mishra, & Cordes, 2019)

Studies in the field of registration are generally divided into two categories: learning-based and non-learning based. In the non-learning category, extensive work has been done in the field of 3D medical image registration (Ashburner, 2007; Avants, Epstein, Grossman, & Gee, 2008; Bajcsy & Kovačič, 1989; Dalca, Bobu, Rost, & Golland, 2016; Sokooti, Saygili, Glocker, Lelieveldt, & Staring, 2016; Thirion, 1995). Some models are based on optimizing the field space of displacement vectors, which include elastic models (Bajcsy & Kovačič, 1989; Kybic & Unser, 2003), statistical parametric mapping (Ashburner, Andersson, & Friston, 2000), free-form deformations with b-spline (Ashburner et al., 2000), and demons(Thirion, 1995). Common formulations include Large Diffeomorphic Distance Metric Mapping (LDDMM) (Ceritoglu et al., 2010; Risser et al., 2011), DARTEL (Ashburner, 2007) and standard symmetric normalization (SyN) (Avants et al., 2008). There are several recent articles in learning-based studies that have suggested neural networks for registering medical images, and most of them require ground truth data or any additional information such as segmentation results (Krebs et al., 2017; Rohé, Datar, Heimann, Sermesant, & Pennec, 2017; Sokooti et al., 2017; Yang, Bian, Yu, Ni, & Heng, 2017)

To the best of our knowledge, no prior study utilized AI for motion correction in the spinal cord fMRI. This study aimed to train a deep learning-based CNN via unsupervised learning to detect and correct motions in axial T2*-weighted spinal cord data. We hypothesize that our method can improve the outcome of motion correction and reduces the preprocessing time as compared to the existing methods.

## 2. Material and Methods

### 2.1 Methods

#### 2.1.1. Unsupervised deep learning network architecture

##### Convolutional Neural Network (CNN) architecture

Assume *M* and *F* are two images of the same slice defined in the *N-dimensional* spatial domain ***Ω*** ⊂ *R*^*N*^. We are focusing on *N = 2* because the type of data we are using is “functional,” containing single-channel grayscale data. Additionally, our network focuses on the Axial view. The fixed image *F* is the reference volume, so it can be the first, middle, average, or any of the volumes, and *M* is the rest of the time-series images. Before training the network, we align *F* and *M* using our fixing Centerline method, which we describe in the following section, so that the only misalignment between the volumes is nonlinear. We then use a convolutional neural network structure similar to UNet (Isola, Zhu, Zhou, & Efros, 2017; Ronneberger, Fischer, & Brox, 2015) to model a *N*_*θ*_(*F,M*) = ∅ function, which includes an encoder and decoder with skip connection (Figure 1): where ∅ is the register map between the two input images and the *θ* learned parameter of the network. In this map, for each voxel *p* ***∈*** *Ω*, there is a position where *F(p)* and the warped image *M*(∅(*p*)) have the same anatomical position. Therefore, our network takes the concatenated images *F* and *M* as input and calculates the registration flow field based on *θ*. In the next step, it uses the spatial transformation operator to warp the moving image based on the flow field and evaluates the similarity between *M* and *F* and *θ* update. Figure 3 shows our introduced architecture and an integrated input by concatenating *F* and *M* in two channels of the 2D image.

**Figure 1.**
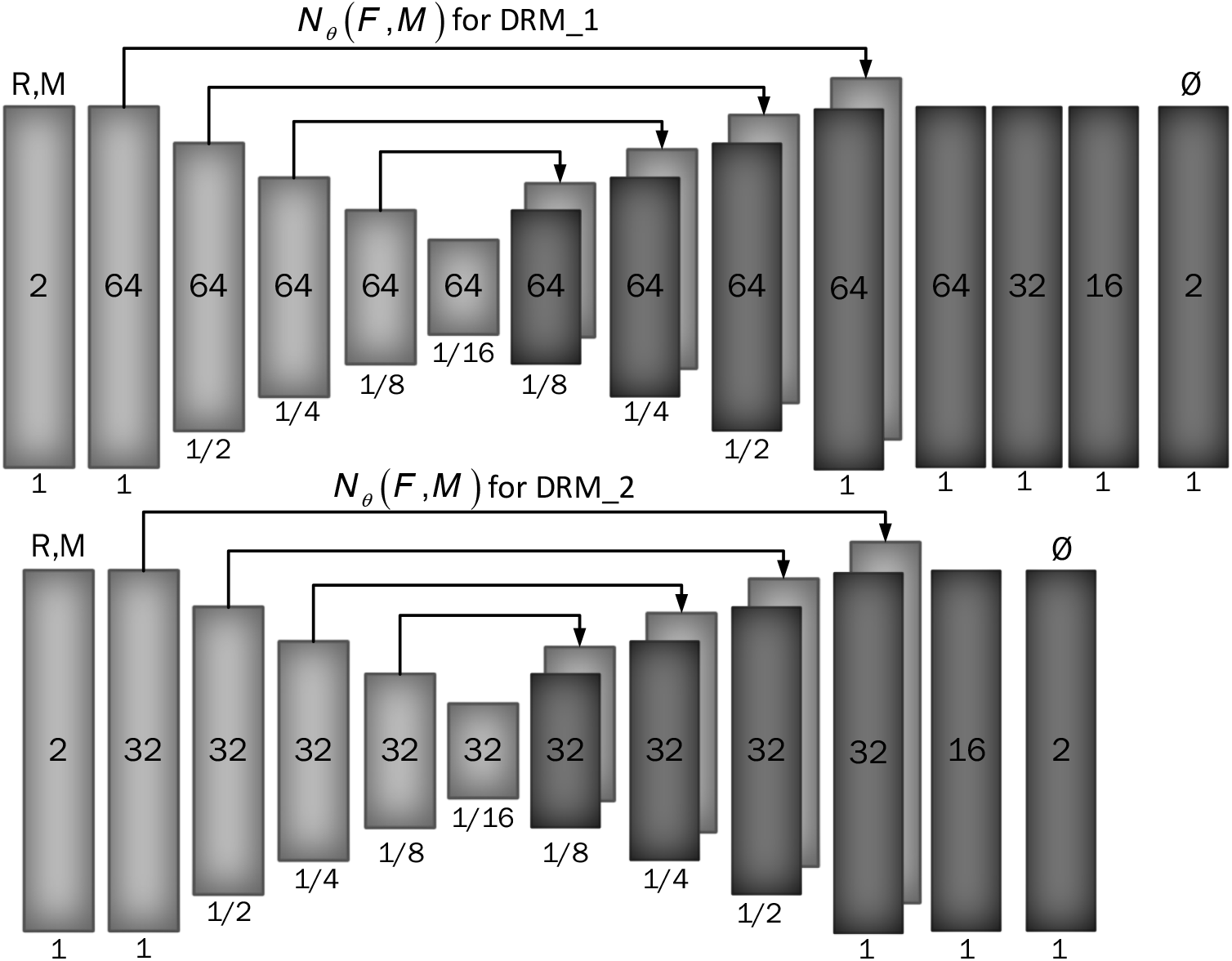
Proposed convolutional architectures implementing g_θ_(F,M). Each rectangle shows a 2D volume in which two fixed and moving images are connected. The number of channels inside each rectangle is shown and the spatial resolution is printed below it according to the input volume. The first model has a larger architecture and more channels than the second model.

In both the encoder and decoder stages, we use two-dimensional convolution with a *3×3* kernel size and leaky Relu activation. The hierarchical properties of the concatenated image pair are captured by the convolution layer, which is required to estimate ∅. We also use stride convolution to decrease the spatial dimensions and get to the smallest layer. During the encoding steps, features are extracted by down sampling, and during the decoding and up sampling steps, the network propagates the trained features from the previous step directly to the layer that generates the registry by using a skip connection. A decoder’s output size (∅) is equal to the input image *M*.

We used two architectures to examine a trade-off between speed and accuracy. These two structures are DRM_1 and DRM_2, which differ in size at the end of the decoder. DRM_1 uses more layers at the end of the decoder, and more channels are used throughout the model (Figure 1).

To find the optimal theta parameter, we used the stochastic gradient descent method to minimize the loss function *ℒ*:

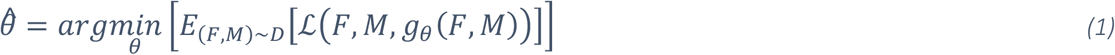

Where *D* is the data scatter. It should be noted that we do not need supervisor information such as Atlas or T1 images.

The *ℒ*_*Unsupervised*_ consists of two parts: *ℒ*_*sim*_, which measures the similarity between *F* and *M*(∅), and *ℒ*_*reg*_, which measures the smoothness of the registration field. So, our total loss function is as follows:

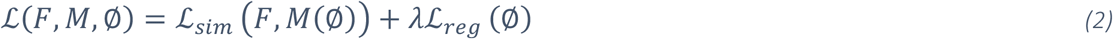

And *λ* is the regulation parameter.

We used two different cost functions for *ℒ*_*sim*_ : mean square error and normalize cross-correlation, which is a common metric due to robust intensity variations. The first cost function, the mean square error is as follows:

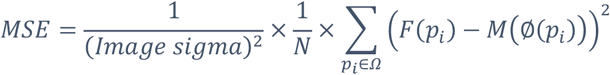

Here *p*_*i*_ is the position of the pixels and Image sigma is equal to one in this work. Also, the fact that MSE is close to zero indicates better alignment. The second cost function, normalizing cross-correlation, is as follows:

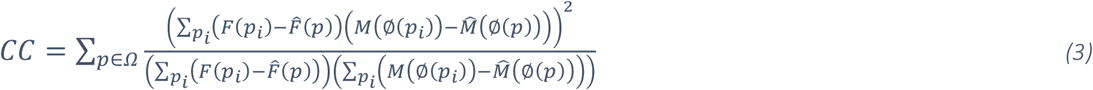

Let *F*(*p*_*i*_)and *M*(∅(*p*_*i*_)) be the image intensities of fixed and moving images respectively, and 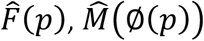 be the local mean at position p, respectively. The local mean is computed over a local *n*^2^ window centered at each position p with *n=9* in this work. Therefore, *p*_*i*_ represent the position within *n*^3^ local windows centered at *p*.

By minimizing *ℒ*_*sim*_, we seek to approximate *M*(∅(*p*)) from *F*(*p*), but it may cause a discontinuity in ∅, so we used Spatial gradients to regulate the deformation field between the voxel’s neighborhood, as follows:

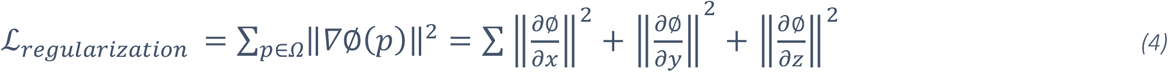

This cost function is applied to the network’s output vectors and controls the size of the vectors by deriving the vectors in each direction.

##### Spatial Transformation Function

The Spatial Transformation Function method learns the optimal parameter by minimizing the difference between the warped image and the reference image. In this way it learns the transformation from the training network and creates a sample grid by the predicted transformation parameters *θ*, which is a set of points that samples the input image to give the output image (Jaderberg, Simonyan, Zisserman, & Kavukcuoglu, 2015). Therefore, to perform a spatial transformation on the input image, the Spatial Transformation Function sampler must take a number of sample points of moving image *M* to produce the transformed image *M*(∅). Also, because the *M*(∅)image is only defined in integer locations, we perform bilinear interpolation in 8 neighboring voxels for the sampler.

#### 2.1.2. Fixing centerline as preprocessing

Before predicting with the network, we align the data in each slice over time using a centerline in the spinal cord. We used the spinal cord toolbox to extract the centerlines, but we had to use interpolation to adjust the points because some of them were outside the anticipated range or were missing. We organized the centerline coordinates by utilizing third-degree b-spline interpolation after identifying the outliers using the interquartile range approach to discover the lower and upper boundaries of the centerline coordinates.

In this step, data is only corrected in 2D, *x* and *y*, and we can choose one or both dimensions (*x, y*, both) for alignment in this approach (Figure **2**). we chose to only correct along the y-direction due to two reasons: 1) we observed that most of the movements in the Axial view of spinal cord data are in the y-direction, also consistent with more distortions along the phase encoding direction. 2) using numerical analysis of the coordinates of the center lines in the x and y directions based on the specific reference, we reached the average variance of 0.52 for x and 1.1 for y, so the scattering of the middle coordinates for the x-direction is about twice less than the y-direction.

**Figure 2.**
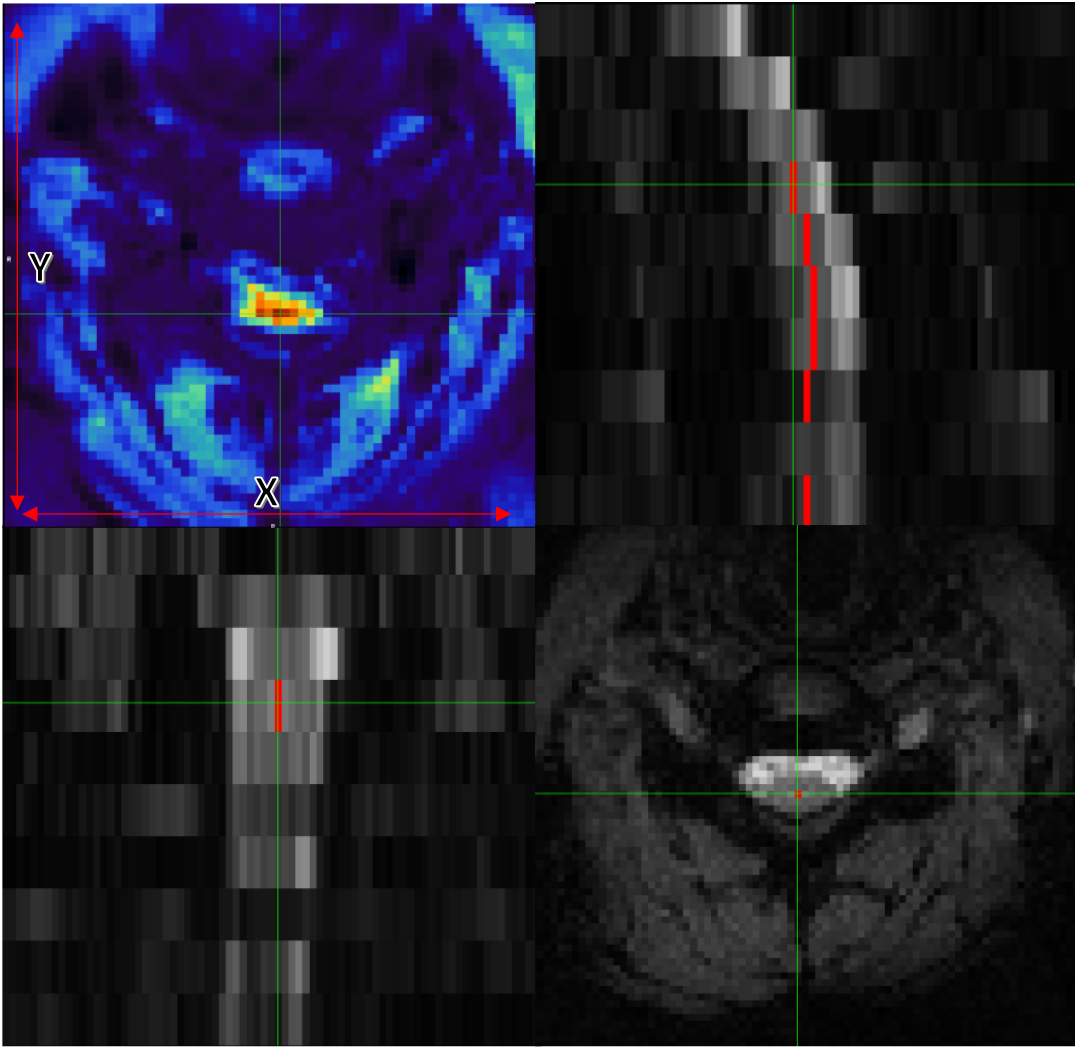
Axial (bottom right), Coronal (bottom left), and Sagittal (top right) views of data with the centerline. The image on the top left also shows tSNR with the x and y guidelines.

**Figure 3.**
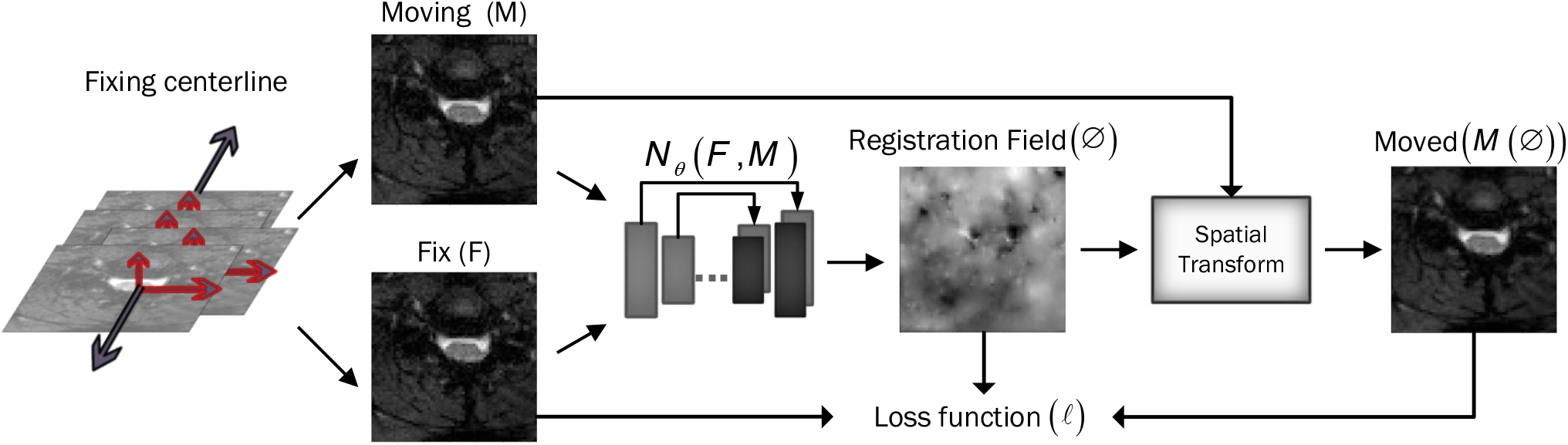
Overview of DeepRetroMoco. As a preprocess, we align the data in two dimensions based on the centerline, and then we register the moving image (*M*) to the fixed image (*F*) by learning function parameters (*N*_*θ*_). During training, an ST was used to warp the moving image with the registration field, and in this operation, the loss function compares M(∅)and *F* using the smoothness of ∅.

### 2.1. Experiments

#### 2.2.1. Dataset

The data used for this experiment includes 30 subjects with T2*-weighted MRI scans acquired from 3T TIM Trio Siemens scanner (Siemens Healthcare, Erlangen, Germany) equipped with a 32-channel head coil and a 4-channel neck coil was used for the imaging to investigate the functional activity in the brain and the spinal cord (Khatibi et al., 2022). All subjects were scanned twice. Five runs were collected in the first session and three runs in the second session. Sessions were acquired one week apart. This resulted in 240 runs. We only used the data from the neck coil and cervical spinal cord in this study.

The dataset included 8–10 slices that covered the cervical spinal cord from C3 to T1 spinal segmental levels and were orientated parallel to the spinal cord at the C6 level. The FoV of the slices was 132 × 132 *mm*^2^, with voxel sizes of 1.2 × 1.2 × 5 *mm*^3^ and a 4 mm gap between them. The flip angle was 90 °, and the bandwidth per pixel was 1263 Hz, resulting in an echo spacing of 0.90 ms. 7/8 partial Fourier and parallel imaging (R = 2, 48 reference lines) was utilized again, resulting in a 43.3 ms echo train length and a 33 ms echo time. Finally, the TR for all slices was 3140 ms, with the exception of three subjects, who had TRs of 3050 ms or 3200 ms (depending on each participant’s coverage within the field of view). We eliminated the lost data due to low quality and differences in data points when compared to other data, and selected data from 27 subjects and 216 runs: 135 runs for training and a total of 81 runs for testing datasets. For training and validation, the training data was divided by 70 to 30%. The data from the validation part was utilized to select and assess the proposed models.

#### 2.2.2. Evaluation

Since there is no gold standard for direct evaluation of functional registration or motion correction performance, we used two functional measures to check the signal strength of each subject or to examine signal variations in the group of volumes after predicting them by the network.

##### 1-temporal signal to noise ratio (tSNR)

Temporal SNR (tSNR) is used to quantify the stability of the BOLD signal time series and is calculated by dividing the mean signal by the standard deviation of the signal over time.

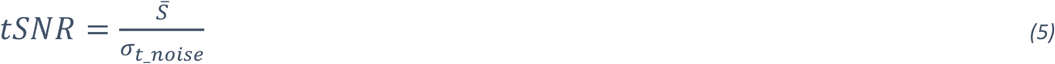

where 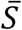 is the mean signal over time and *σ* is the standard deviation across time. A better motion correction algorithm will result in greater tSNR values by reducing signal variations in the BOLD time-series due to motion.

##### 2-DVARS

DVARS (D, temporal derivative of time courses, VARS, variance over voxels) shows the signal rate changes in each fMRI data frame. In an ideal data series, its value depends on the temporal standard deviation and temporal autocorrelation of the data (Nichols, 2017) and calculates the changes in the values of each voxel at each time point compared to its previous time point (Power, Barnes, Snyder, Schlaggar, & Petersen, 2012). DVARS was calculated in the whole image to find a metric that demonstrated the standard deviation of temporal difference images in the 4D raw data (Power et al., 2014). DVARS was calculated using the following equation:

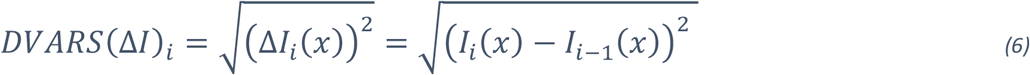

In this equation, ΔI_*i*_(*x*) is used as local image intensity on the frame. DVARS could result in more accurate modeling of the temporal correlation and standardization because it is obtained by the most short-scale changes (Nichols, 2017). The best value for this parameter is zero, and the closer it is to zero, the better the result.

We extracted the tSNR and DVARS parameters of output results by using the SCT toolbox and the FSL toolbox (Woolrich et al., 2009). For more accurate analysis of the tSNR parameter, we manually segmented the data into two parts, spinal cord and CSF, using the FSLeyes toolbox. Analyses compared the outcome of SCT and our method (DeepRetroMoco).

#### 2.2.3. Statistical Analysis

All statistical analyses were carried out using IBM SPSS Statistics (V. 25 IBM Corp., Armonk, NY, USA) with α < 0.05 as the statistical significance threshold. The Kolmogorov-Smirnov test was used to determine the normality of the parameters. For statistically significant results, the mean of normal data for each method was processed using one-way ANOVA with repeated measures in within-subjects comparison, followed by a multiple comparison post-hoc test with Bonferroni correction.

#### 2.2.4. Implementation

In our experiment, we trained our deep learning network with and without using Fixing Centerline as preprocessing for the network. The number of our training epochs is 200 with 150 iterations for each epoch. We used Keras with TensorFlow backend (Abadi et al., 2016) on NVIDIA GEFORCE RTX 1080 to train our network, and it took an average of 23 hours to run the network. We also used Google Colab to review models and learn various parameters, which we worked on much faster.

The optimization parameter we used was Adam, with a learning rate of 1*e*^−4^ (Kingma & Ba, 2015). We trained our two simple (DRM_2) and complex (DRM_1) models designed by two different cost functions (NCC, MSE) and different lambda up to convergence time.

In our study, we designed a data generator to deliver fMRI data to the network. The data generator randomly selects the subject and slice and then selects a pair (fixed and moving) from the corresponding volume to the size of the data batch.

In comparison between the models, we chose the model that has better results in terms of our desired metric (tSNR) on the validation data. Then, we select one of the cost functions. Our code and model parameters are available online at https://github.com/mahdimplus/DeepRetroMoco

## 3. Result

### 3.1 Model selection

Table 1 displays the average of our method’s tSNR values in the validation data utilizing two distinct cost functions. The first model, DRM 1, outperforms DRM 2 in both Losses MSE and NCC by a slight margin. Furthermore, when the validation data of two cost functions in the first model are examined, NCC with an average of 10.13 ± 1, has better outcomes for the motion correction target based on the tSNR and statistical analysis, t(39) = 2.63, p<0.05.

**Table 1.** Average tSNR for two types of our model, DRM-1 and DRM-2. Standard deviations are in parentheses. The averages are computed over all validation data. And in both models, regardless of the type of cost function, the first model is selected. (df=39)

| **Model** | **Loss type** | **Mean. tSNR (std)** | **F-value** | **P-value** |
| --- | --- | --- | --- | --- |
| DRM_1 | MSE | 6.18(0.6) | 12.408 | <0.001 |
| DRM_2 |  | 5.96(0.7) |  |  |
| DRM_1 | NCC | 10.13(1) | 2.632 | <0.05 |
| DRM_2 |  | 10.07(1.3) |  |  |

### 3.2 Test statistic report

A repeating one-way measurement ANOVA was used to compare the influence of motion correction techniques on test data in sct_fmri_moco (De Leener et al., 2017) and DeepRetroMoco, a deep neural network-based motion correction tool.

In a statistical comparison of tSNR parameters in the spinal cord, this parameter increased significantly from 7.104 ± 2.41 to 16.072 ± 3.09 arbitrary units (AU) (Table 3). Mauchly’s Test of Sphericity revealed that the assumption of sphericity had been violated, χ^2^(9) = 2.324, p < .313, and thus a Greenhouse-Geisser correction was used. The motion correction algorithm had a significant effect on the tSNR parameter in the spinal cord, F (2, 160) = 862.572, p < .0001. Post hoc multiple comparisons using the Bonferroni correction revealed that the DeepRetroMoCo had a significantly higher mean tSNR in the spinal cord than the other motion correction method and raw data (p<.0001). Figure 4 depicts the significant difference between the groups using a violin plot.

**Figure 4.**
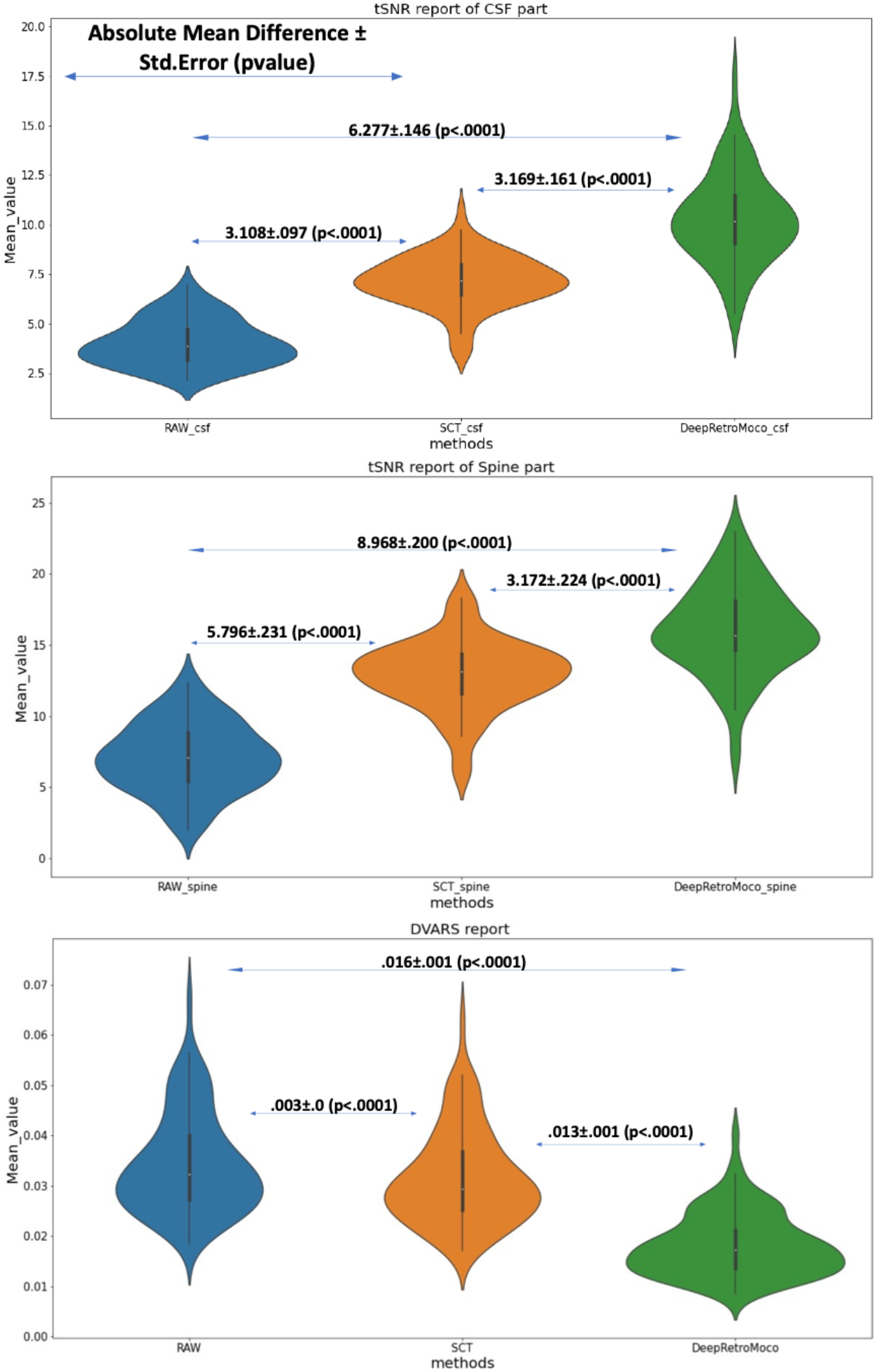
This figure depicts the mean and standard deviation of the SNR on the Spine and CSF sections (two top figures) that were manually segmented, as well as DVARS (bottom figure) with three types of results. RAW data that hasn’t been corrected, SCT results, and DeepRetroMoco results are the three groups. The absolute mean difference + standard error (p value) between groups is also reported. *. The mean difference is significant at the .05 level.

**Figure 5.**
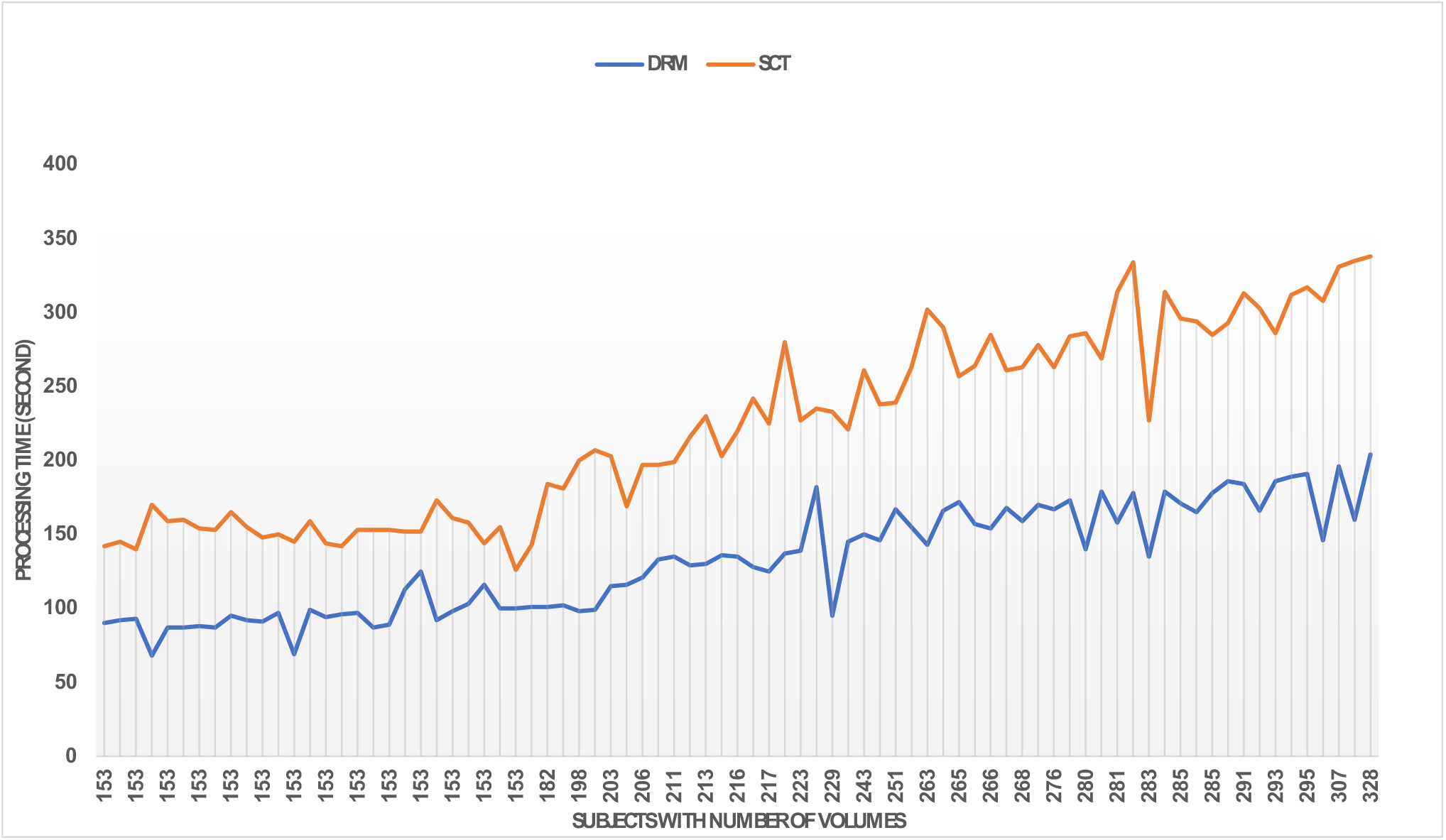
Comparing the speed of the two methods sct_fmri_moco (SCT) and DeepRetroMoco (DRM). Processing time is measured in seconds to correct the motion on all volumes.

**Figure 6.**
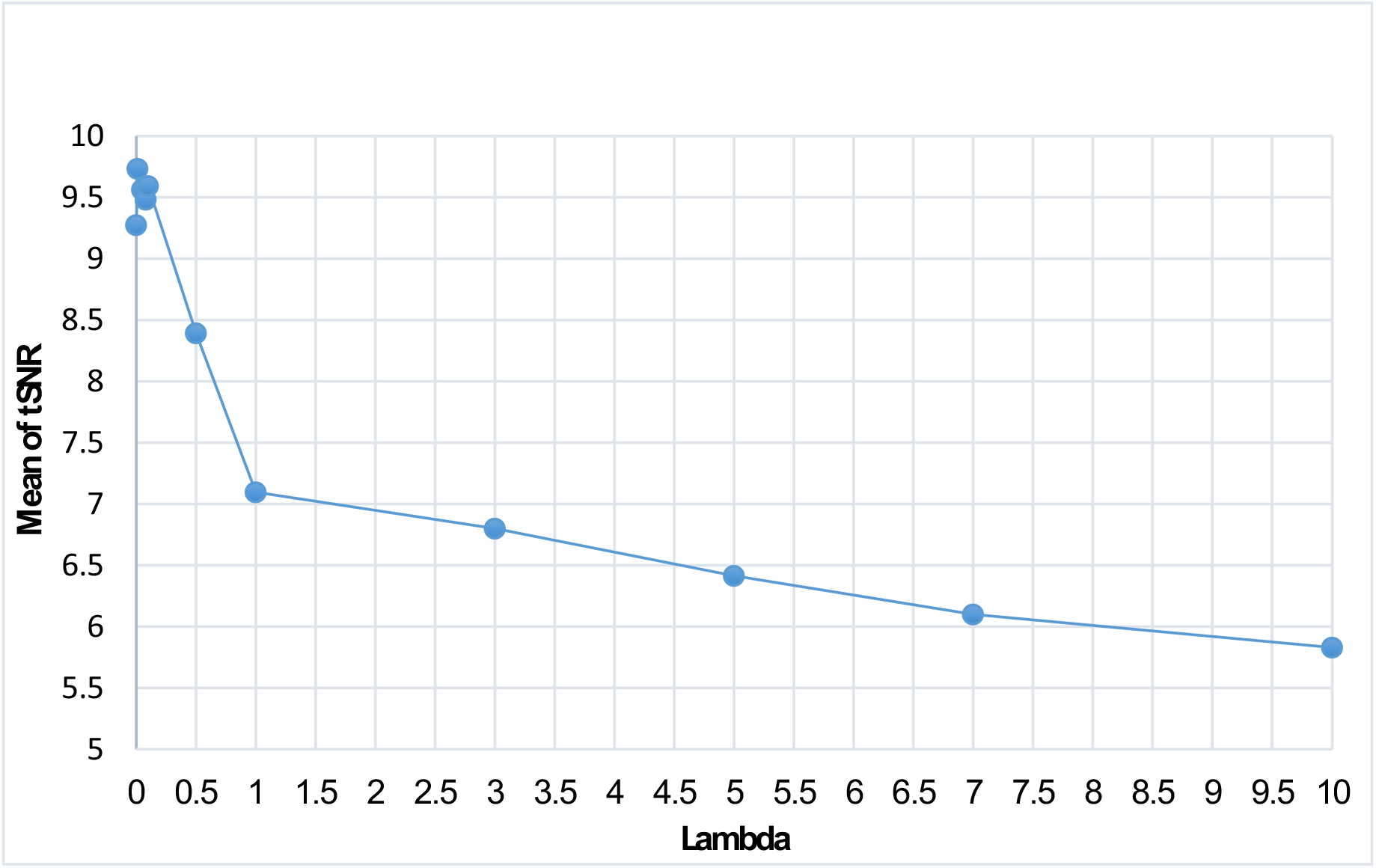
Effect of different λ modes for DRM_1 based on tSNR. Lambda 0.01 has a maximum tSNR and shows the best results.

The tSNR in CSF increased significantly from 4.038 ± 1.17 to 10.315 ± 2.25 arbitrary units (AU) (Table 2 and Table 3). Mauchly’s Test of Sphericity revealed that the sphericity assumption had been violated, χ^2^(9) = 27.772, p < .0001, and thus a Greenhouse-Geisser correction was applied. The motion correction algorithm had a significant effect on the tSNR parameter in CSF F (2, 160) = 949.72, p < .0001. Post hoc multiple comparisons using the Bonferroni correction revealed that the DeepRetroMoCo’s mean tSNR in CSF was significantly higher than the other motion correction method and raw data (p<.0001). Figure 4 depicts the significant difference between the groups using a violin plot.

**Table 2.** Summary of tSNR as an Image quality parameter between different motion correction methods (df=4)

|  |  |  |  |  |
| --- | --- | --- | --- | --- |
| *tSNR* |  | *Mean (std)* | *F value* | *p-value* |
| Spinal cord | Main Image | 7.1043(2.41) | 1004.249 | <0.001 |
|  | sct_fmri_moco | 12.90(2.44) |  |  |
|  | DeepRetroMoCo | 16.072(3.09) |  |  |
| CSF | Main Image | 4.0387(1.17) | 938.842 | <0.001 |
|  | sct_fmri_moco | 7.1469(1.31) |  |  |
|  | DeepRetroMoCo | 10.3156(2.25) |  |  |

**Table 3.** Summary of DVARS as an Image quality parameter between different motion correction methods (df=4)

|  |  |  |  |
| --- | --- | --- | --- |
| DVARS | Mean (std) | F value | p-value |
| Main Image | .0343(.009) | 176.446 | <0.001 |
| sct_fmri_moco | .0316(.009) |  |  |
| DeepRetroMoCo | .0182(.006) |  |  |

DVARS decreased statistically significantly from 0.034 ± 0.009 to 0.182± 0.006 arbitrary units (AU) (Table 4). Mauchly’s Test of Sphericity revealed that the sphericity assumption had been violated, χ^2^(9) = 64.966, p < .0001, and thus a Greenhouse-Geisser correction was applied. The motion correction algorithm had a significant effect on the DVARS parameter, F(2, 160) = 309.349, p < .0001. Post hoc multiple comparisons using the Bonferroni correction revealed that the DeepRetroMoCo had significantly lower DVARS than the other motion correction methods and raw data (p<.0001). Figure 4 depicts the significant difference between the groups using a violin plot.

**Table 4.**
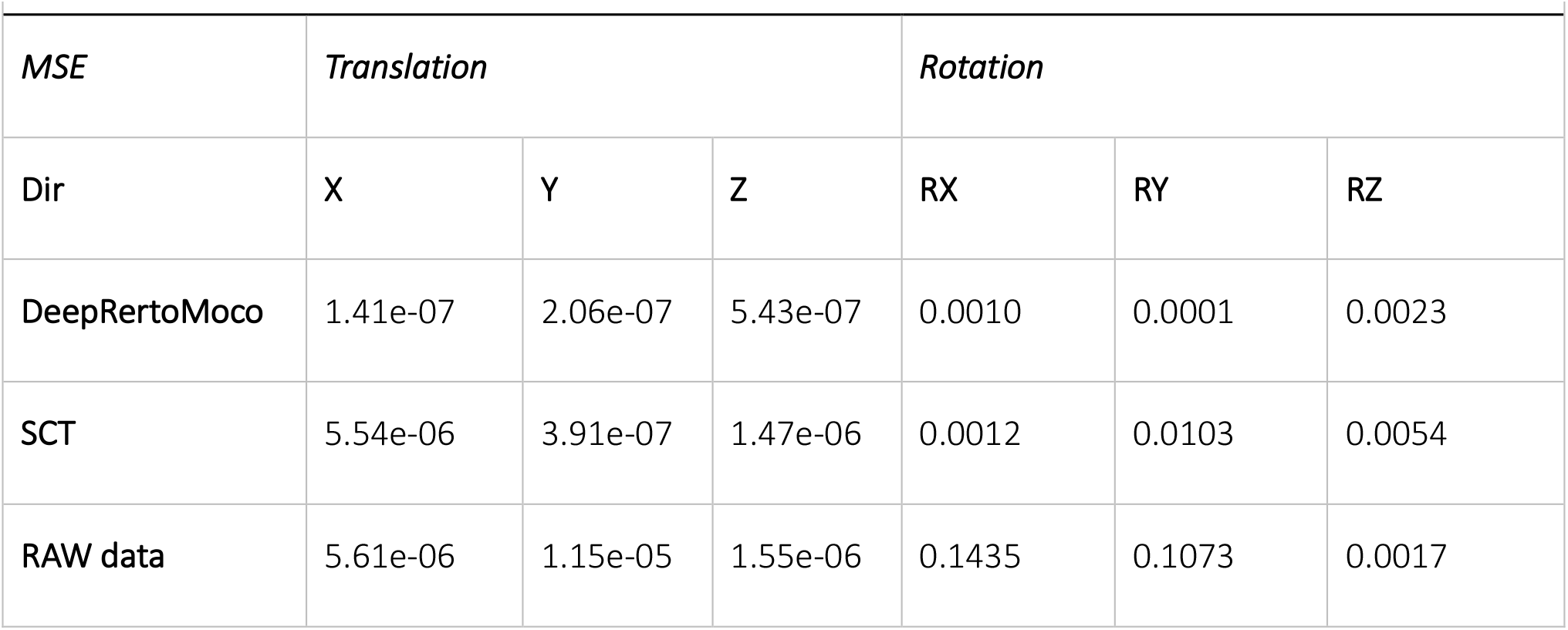
Mean square error of 3 groups of our data in 6 directions such as Translation in X, Y, Z and Rotation in X, Y, and Z directions.

### 3.3 Compare with other methods

Further, we used the FSL gold standard method and MC_FLIRT for the estimation of movements in three groups of our data. Group one contains the RAW data which is not corrected for motion. Group two includes the result of the SCT toolbox, sct_fmri_moco and the third group contains the DeepRetroMoCo results. The reference volume was set to the first volume and the DOF was 6.

We used the MSE parameter to analyze the results, setting the reference line for actual movement to zero and the predicted movement to the numbers reported by FSL. As reported in Table 5, the raw data had the most movement in all directions, followed by the SCT and DeepRetroMoCo results, which had the most movement respectively.

### 3.4 Processing speed

The implementation and calculation are carried out in a workstation with Intel® Core (TM) i7-4720HQ CPU at 2.60Hz and 16.0GB memory. No explicit parallelization was implemented in the Python script. The computation time of the motion correction procedure in sct_fmri_moco and DeepRetroMoco changes with the number of volumes of fMRI raw data. Average computation times (± Stdv) were 222.54±63.64 seconds and 131.91±35.94 seconds for sct_fmri_moco and DeepRetroMoco respectively and demonstrates a significant reduction of ∼40.72% in computation time. This operation for SCT contains the slice-by-slice registration plus regularization across the Z, and for DeepRetroMoCo contains fixing centerline plus registration via a network.

### 3.5 Regularization Analysis

With different lambda parameters, we examined the mean tSNR for the test data. With the NCC cost function, the optimal tSNR for model 1 occurred when lambda was 0.01. In this section, the mean tSNR is applied to the entire spinal cord; lambda=0 indicates no regularization. As shown, the results deteriorate dramatically as the regularization term is increased. As a result, lambda’s actions do not help to improve performance and may have a negative impact on the results for the NCC cost function and the first model, which is more complex.

### 3.6 Correlation Coefficient Analysis

We further calculated the Pearson correlation coefficient (CC) between the corrected and reference volumes to examine the similarity of the two images in a linear fashion. For both the raw data and the motion-corrected data, we calculated the average CC per subject. Since linear relationships between variables are preserved by a linear transformation, the correlation between x and y would remain unchanged following a linear translation (Carroll et al., 1985). We found a large average CC value in our algorithm (0.90±0.02) compared to the raw data (0.70±0.17), which shows that our trained network (DeepRetroMoco) uses a significant proportion of linear transformations due to our rigorous regularization.

## 4. Discussion

Since the spinal column’s voluntary and non-voluntary movements lead to non-optimal shimming, the effects of motion artefacts cannot be fully eliminated even after perfect conventional retrospective motion correction of successive functional volumes in image space (Eippert, Kong, Jenkinson, Tracey, & Brooks, 2017). If spinal column movements are small, motion correction is a useful step to improve the data quality for subsequent statistical data analysis. Our findings demonstrate that deep learning-based motion correction, DeepRetroMoco improves the quality of spinal cord fMRI data acquired in the axial field of view that has an effect on the pre-processing step. These improvements are at least in part due to improved tSNR and DVARS parameters compared to conventional algorithms introduced in the SCT data processing toolbox.

Instead, here we aimed to use a deep learning-based method potential to decrease preprocessing step for spinal cord fMRI data that strongly affected by motion. We found significant differences in the time of processing to implement DeepRetroMoco compared to the sct_fmri_moco algorithm.

As previously mentioned, the majority of leaning-based methodologies require additional data or ground truth. We don’t need this information, which is another clear distinction between our approach and earlier research. The previous two works (Li & Fan, 2017; Vos, Berendsen, Viergever, Staring, & Išgum, 2017) reported unsupervised methods that are close to ours. Both use the CNN neural network with spatial transformation function (Jaderberg et al., 2015), which warps images on top of each other and has significant problems: they only operate on a limited subset of volumes and only support small transformations. In addition a recent study (Balakrishnan, Zhao, Sabuncu, Dalca, & Guttag, 2018) and our network improved the problems mentioned and helped to solve them by designing a satisfactory model in the spinal cord data. Other methods (Vos et al., 2017) use regularization that is determined only by interpolation methods.

The DeepRetroMoco replaces a costly optimization problem for each image pair, with a function optimization that is collected over a data set during a training step. This notion could be replaced with previous motion correction algorithms, especially on spinal cord data which traditionally relies on complex, non-learning-based optimization algorithms for each input. Although implementing this network requires a one-time network training on a single NVIDIA TITAN X GPU with training data, it takes less than a second to register a pair of images. Due to the growing need for medical images for further investigation in less time, our solution, which is a learning-based method, is preferable to non-learning-based methods.

## 5. Limitation and Future Works

The acquisition of spinal cord fMRI data is made in two ways: GRE-EPI acquisition sequence in axial and FSE or SE-HASTE acquisition sequence in sagittal field of view. The field of view and data-set orientation were axial in this study, and all motion correction methods and preprocessing steps were performed specifically on axially oriented data in the cervical spine; however, some studies performed spinal cord fMRI acquisition in the sagittal orientation. Respiration and heartbeat cause changes in CSF flow around the spinal cord, which results in magnetic susceptibility during the GRE-EPI image acquisition and preprocessing steps.

Furthermore, we had access to two variables during this method: the centerline reference and the fixed image reference. It was set to the first volume in our network. We discovered that the proper selection of these two parameters could have a significant impact on the final results. Because our network is flexible enough to accept any reference, including first, mean, middle, and any other desired volume, we propose that the best reference for each data be selected by designing the appropriate method for future work.

## 6. Conclusion

Owing to the bulk and physiological motion corrupted spinal cord fMRI data, the statistical significance of the activation maps decreases, and the likelihood of false activations increases. As a result, a motion correction algorithm is required for acceptable single and group fMRI data analysis. In this study, we proposed DeepRetroMoco, an unsupervised learning-based approach based on advanced CNN models, that requires no supervised information such as ground truth registration fields or anatomical landmarks. Additionally, when compared to conventional methods, the use of DeepRetroMoco motion correction in spinal cord fMRI appears to be impressive in terms of increasing tSNR, reducing false positives, and increasing sensitivity, especially in cases of significant spinal cord motion. Furthermore, the statistical evaluation of DVARS as an fMRI quality measure, as well as the time of implementation on a cervical spinal cord fMRI dataset, demonstrated the superiority of the proposed framework in our experimental study and also is an easy-to-integrate tool for more accurate and faster motion correction for denoising in spinal cord fMRI applications.

